# Fucosyltransferase 4 shapes oncogenic glycoproteome to drive metastasis of lung adenocarcinoma

**DOI:** 10.1101/752519

**Authors:** Hsuan-Hsuan Lu, Shu-Yung Lin, Rueyhung Roc Weng, Yi-Hsiu Juan, Yen-Wei Chen, Hsin-Han Hou, Zheng-Ci Hung, Giovanni Audrey Oswita, Yi-Jhen Huang, Jin-Yuan Shih, Chong-Jen Yu, Hsing-Chen Tsai

**Author notes:** Contributed equally (Chong-Jen Yu, Hsing-Chen Tsai). Correspondence: Hsing-Chen Tsai, MD, PhD., Graduate Institute of Toxicology, College of Medicine, National Taiwan University, No.1 Jen Ai Road Section 1, Rm1424, Taipei, Taiwan 10051, #88797 or Chong-Jen Yu, MD, PhD.; Department of Internal Medicine, National Taiwan University Hospital, No. 7, Zhongshan S Rd, Zhongzheng District, Taipei City, 10002, Taiwan. #62905.

## Abstract

Aberrant fucosylation plays a critical role in lung cancer progression. Identification of the key fucosyltransferase as a therapeutic target may refine lung cancer management. Here, we identified a terminal α1,3-fucosyltransferase, FUT4, as the key prognostic predictor for lung adenocarcinoma through transcriptomic screens in lung cancer cohorts. Overexpression of FUT4 promotes lung cancer invasion, migration and cell adhesion *in vitro* and provokes distant metastases in mouse xenograft models. RNA-seq and glycoproteomics analyses revealed that FUT4 mediates aberrant fucosylation of intracellular transport and signal transduction proteins, which facilitates concurrent transcriptional activation of multiple cellular processes, including membrane trafficking, cell cycle, and major oncogenic signaling pathways. Notably, knockdown of FUT4 markedly curtailed lung colonization and distant metastases of lung cancer cells in mouse xenograft models. In addition, the metastatic phenotype provoked by FUT4-mediated fucosylproteomic networks can be diminished by targeted pathway inhibitors. Collectively, FUT4 represents a promising therapeutic target in lung cancer metastasis. Our data highlight the potentials for integration of glycomics into precision medicine-based therapeutics.

## Introduction

Non-small cell lung cancer is a disease with genetic and gene regulatory complexities. Molecular analysis of individual tumors is essential for clinical choice of therapeutic strategies in the era of precision medicine. Despite the growing dimensions of lung cancer-associated molecular abnormalities, current practices for lung cancer therapy rely primarily on pathological, mutational, and immunological characterizations. Abundant evidence indicates that aberrant glycosylation plays critical roles in fundamental steps of tumor development and progression, including cell-cell/cell-matrix interactions, metastasis, cancer metabolism as well as immune surveillance^1–3^. In particular, patterns of fucosylation, which adds a fucose (6-deoxy-L-galactose) residue to surface oligosaccharides/proteins catalyzed by the fucosyltransferase (FUT) gene family, are frequently altered in various types of cancer^4, 5^. Depending on the site of oligosaccharide chain to which the fucose is added, two types of fucosylation— core and terminal fucosylation—were defined^4^. So far, there are 13 FUTs identified in human, each of which catalyzes the synthesis of fucosylated glycans with designated glycosidic linkages and targets different substrate proteins/glycans in a tissue-specific manner.

In search for the key fucosyltransferases underlying aberrant fucosylation patterns specific for lung cancer progression, we and others have previously discovered that fucosyltransferase 8 (FUT8), the only enzyme responsible for the core fucosylation with α1,6-linkage, mediated the malignant phenotypes of non-small cell lung cancer^6, 7^. On the other hand, clinical significance of fucosyltransferases involved in terminal fucosylation (α1,3-or α1,4-linkage) in lung cancer remains controversial. Past studies have demonstrated aberrant expression of terminal fucosylated epitopes such as Lewis antigens in non-small cell lung cancer tissues^8–10^. Nevertheless, the prognostic values of various Lewis antigens appeared to differ. Expressions of sialyl Lewis x (sLe^x^) and Lewis x (Le^x^) were associated with shortened survival times^10^. In contrast, Lewis y (Le^y^) seemed to predict better survival or confer limited clinical significance^8, 9^. Furthermore, while some groups found that terminal fucosyltransferases such as FUT4 or FUT7 may promote lung cancer progression^11–13^, another group demonstrated that FUT4-or FUT6-mediated fucosylation of epidermal growth factor receptor (EGFR) could suppress EGFR dimerization and activation^14^. As multiple fucosyltransferases (FUT3-7, 9-11) are involved in terminal fucosylation and Lewis antigen synthesis, a systemic approach on all α1,3-or α1,4-fucosyltransferases with large scale clinical correlation in individual subtypes of non-small cell lung cancers, followed by in-depth mechanistical and molecular studies is needed to identify the key enzyme that accounts for malignant phenotype of lung cancer and decipher the complex molecular networks involved in cancer progression.

In the present study, our group examined 81 surgically-resected tumor tissues from patients with non-small cell lung cancer for molecular and prognostic correlations on all terminal α1,3-or α1,4-fucosyltransferases, and independently validated our results with the TCGA lung cancer cohorts for cross-ethnic generalization. We identified fucosyltransferase 4 (FUT4) as the main indicator for poor clinical outcome in lung adenocarcinoma patients. We conducted mechanistic and functional studies *in vitro* and *in vivo* by altering FUT4 expression levels in lung cancer cells, and deciphered the molecular networks affected by FUT4 through integrated transcriptomic and glycoproteomic analyses. We found that FUT4 activates intracellular transport machinery and enhances oncogenic signaling via aberrant fucosylation of membrane trafficking components or signaling cascade proteins. Finally, we demonstrated therapeutic implications of our findings by directly targeting FUT4 or by FUT4-related pathway inhibitors. Our study pinpointed a clinically-significant fucosylation enzyme for the malignant phenotype of lung adenocarcinoma, and delineated its complex glycoproteomic networks for developing therapeutics for cancer progression in the future.

## Results

### Higher levels of FUT4 is associated with poor prognosis in lung adenocarcinoma

Terminal fucosyltransferases, such as FUT 3-7 and 9-11, were known to catalyze fucosylation with α1,3- and α1,4-linkage. In order to decipher clinicopathological impact of these fucosyltransferases, we examined expression of FUT 3-7 and 9-11 in the genome-wide RNA-seq data of 81 primary lung cancer tissues (adenocarcinoma, n=44; squamous cell carcinoma, n=37) from National Taiwan University Hospital (NTUH) (GEO: GSE120622). We observed high expression of FUT3, 4, 6, 10, 11 in lung cancer tissues, while little expressions of FUT5, 7 and 9 were detected (Fig 1A). Interestingly, similar patterns were observed in another cohort of 517 lung adenocarcinoma and 502 squamous cell patients from the The Cancer Genome Atlas (TCGA) despite the ethnic differences and distinct mutational profiles between the two cohorts (Fig 1A)^15–18^. To determine the clinical significance of individual fucosyltransferases in terms of overall survival, we divided the patient cohorts into high-expressing and low-expressing groups by the median value of individual fucosyltransferases and performed a survival analysis on the two groups using Cox proportional hazard model (Fig 1B). The analysis in lung adenocarcinoma patients demonstrated a negative prognostic effect of FUT4 (*, hazard ratio 2.172 ± 0.217; *P* = 0.0003, Cox regression model), and protective effects of FUT1 (†, hazard ratio = 0.497 ± 0.220, *P* = 0.0015) and FUT10 (†, hazard ratio = .649 ± 0.217, *P* = 0.0047) on overall survivals (Fig 1B). Kaplan-Meier curves also showed significantly worse overall survival of the FUT4^High^ group (*P* = 0.00068; Fig1C). Notably, we did not observe a significant association between FUT4 expression and overall survival in lung squamous cell carcinoma (*P* = 0.23; Fig 1D and Table S1), indicating a distinct glycobiological mechanism may be involved in the progression of this cancer subtype. Moreover, to investigate potential functional coordination between different FUTs, we performed correlation analysis based on mRNA levels of FUTs in 517 lung adenocarcinoma tissues. We found little or modest concordance between the expressions of FUT4 and other α1,3- or α1,4-fucosyltransferases (Pearson correlation coefficient between −0.1 to 0.22, Fig 1E). This suggests the unique role of FUT4 plays in promoting the aggressive phenotypes of lung adenocarcinoma. In contrast, while not having prognostic values, expressions of FUT3 and FUT6 showed a strong correlation (Pearson correlation coefficient, R = 0.73; Fig 1E), in consistent with a previous study showing a functional coordination between these two enzymes in colorectal cancers^19^. Moreover, expression levels of FUT4 in 107 surgically-removed lung adenocarcinoma tissues at NTUH were significantly upregulated compared the adjacent normal lung tissues measured by quantitative real-time PCR (*p*=0.001) (Fig 1F), indicating that elevated FUT4 expression may contribute to the tumorigenesis of lung adenocarcinoma.

**Figure 1.**
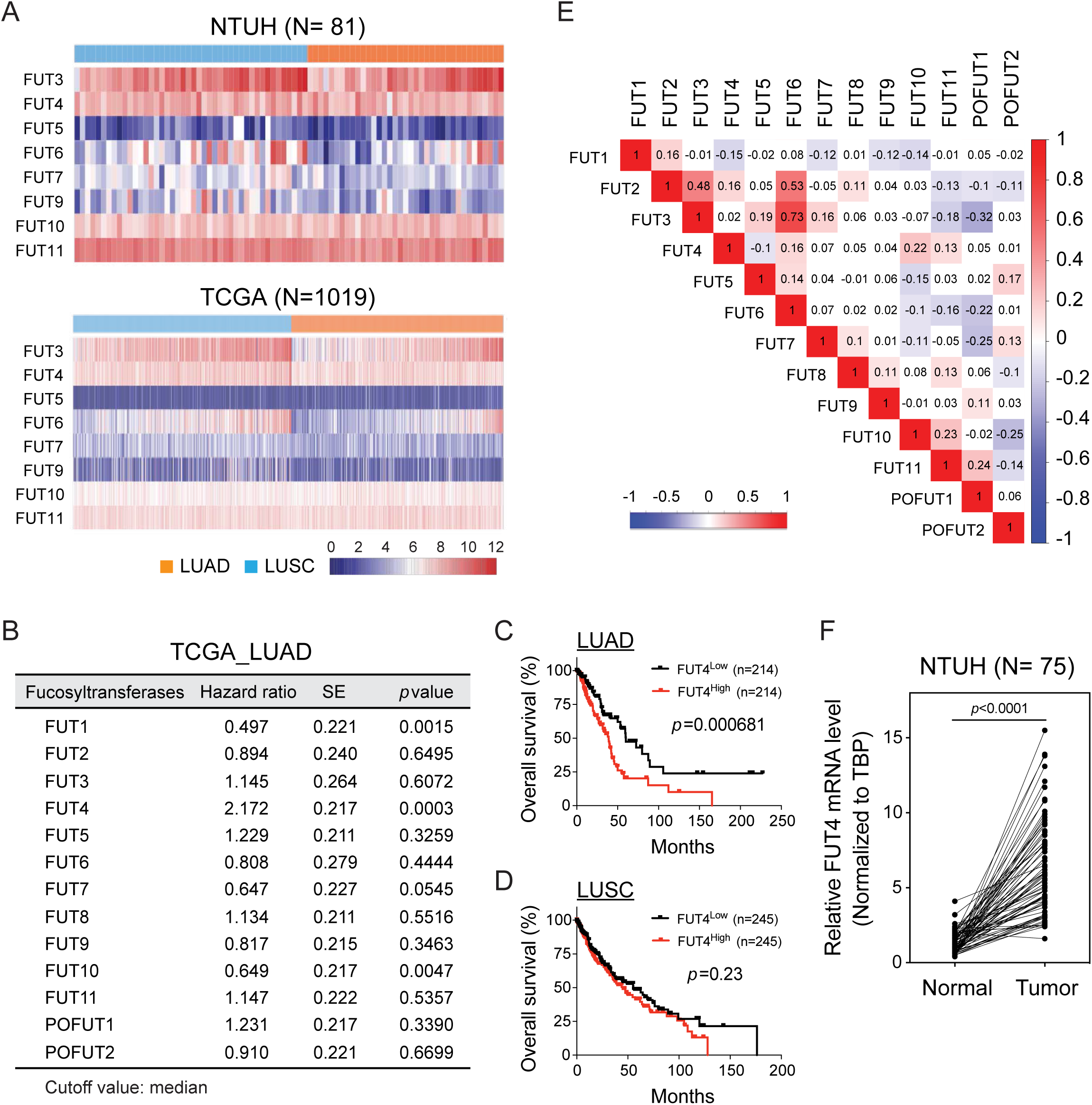
High FUT4 expression in lung adenocarcinoma (LUAD) is associated with poor prognosis. A. Heatmap of RNA-seq transcriptomic analysis for eight α-(1,3)-fucosyltransferases in the surgically-resected lung cancer tissues at National Taiwan University Hospital (LUAD, N=44 and LUSC, N=37) and from TCGA database (LUAD, N=517 and LUSC, N=502), respectively. LUAD: lung adenocarcinoma. LUSC: lung squamous cell carcinoma. B. Relationships between expression levels of individual fucosyltransferases and overall survival in patients with lung adenocarcinoma. (N=428, *High vs. Low expressions, cutoff=median; Cox proportional hazard models*) C, D. Kaplan-Meier curves of overall survival for patients with lung adenocarcinoma (LUAD) (C) or squamous cell carcinoma (LUSC) (D). Patients were divided into two groups based on the median value of FUT4 expressions. E. The correlation matrix of all members in the FUT family based on the RNA-seq data of the TCGA lung adenocarcinoma samples. Numbers in the colored box represent Pearson correlation coefficients. Red and blue colors denote positive and negative correlations with statistical significance (*p* < 0.05), respectively. F. mRNA levels of FUT4 in lung adenocarcinoma tissues compared to those of paired adjacent normal tissues from National Taiwan University Hospital (N=75). Data information: *p* value in (B, C, D) was calculated by log-rank test, and in (F) was calculated by Wilcoxon matched-pairs signed rank test.

### FUT4 promotes aggressive phenotypes of human lung cancer cells

To decipher the mechanisms of FUT4 that account for poor prognosis in lung cancer patients, we established FUT4-overexpressing lung adenocarcinoma cell line models using the tetracycline-controlled transcriptional regulatory system for a series of *in vitro* and *in vivo* mechanistic studies. First, we examined expression levels of FUT4 in a panel of 17 human non-small cell lung cancer lines. Higher expressions of FUT4 were observed in 13 of the 17 cell lines as compared with immortalized BEAS-2B lung epithelial cells (Fig 2A). Subsequently, we stably transfected two human lung adenocarcinoma cell lines, A549 (Fig 2B) and CL1-0 cells (Fig 2E), with FUT4. For A549 cells, we selected two clones with different levels of FUT4 expression— A549_FUT4^high^ and A549_FUT4^med^ – to investigate the dose-responsive relationship of FUT4 protein. We found that A549_FUT4^high^, A549_FUT4^med^, showed markedly increased invasion abilities through Matrigel^®^-coated Boyden chamber (Fig 2C). Moreover, single cell migration assays in a live-cell imaging and tracking system revealed increased migration abilities of A549_FUT4^high^ and A549_FUT4^med^ cells. (Fig 2D). Interestingly, A549_FUT4^med^ cells showed similar invasion and migration abilities with A549_FUT4^high^ cells, which indicates that moderate elevation of FUT4 protein level was sufficient to cause significant functional changes. Similarly, CL1-0_FUT4 cells showed enhanced invasion ability (Fig 2F) although there was no significant change in cell migration velocity (Fig 2G).

**Figure 2.**
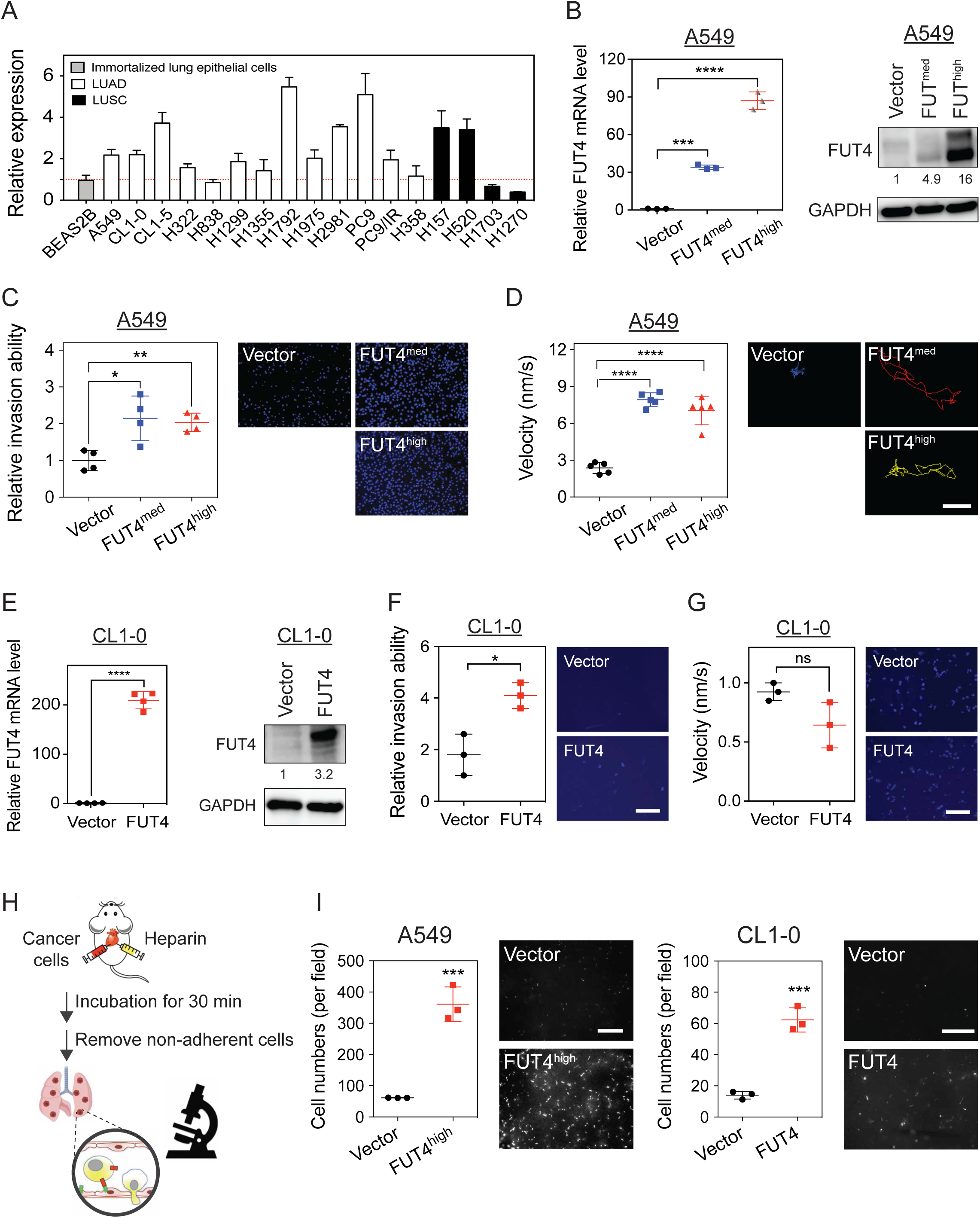
FUT4 promotes aggressive phenotypes of human lung cancer cells. A. FUT4 mRNA expressions in various human lung cancer cell lines analyzed by quantitative realtime PCR. All experiments were performed in triplicates and presented as mean ± standard errors. ADC: adenocarcinoma. SCC: squamous cell carcinoma. B. FUT4 mRNA (*left panel*) and protein levels (*right panel*) in A549 human lung cancer cells transduced with FUT4 (A549_vector, A549_FUT4^med^, A549_FUT4^high^) measured by quantitative real-time PCR and western blots. C. Dot plots showing relative invasion ability of A549 cells measured by matrigel-based transwell invasion assays. Representative images of cells with DAPI nuclear stains in the lower chambers are shown in the *right panels*. Relative invasion ability was calculated using the cell number in the lower chamber of transwell system for each clone compared to that of the vector control. D. Dot plots showing migration velocity of A549 cells measured by single cell tracking assays under fluorescence microscope. Data were analyzed using Metamorph® software. The paths of cell migration were delineated in the *right panels* using pseudo-colors. E. FUT4 mRNA (*left panels*) and protein levels (*right panels*) in CL1-0 human lung cancer cells, which were overexpressed with FUT4 (CL1-0_vector, CL1-0_FUT4) measured by quantitative real-time PCR and western blots. F. Dot plots showing relative invasion ability of CL1-0 cells measured by matrigel-based transwell invasion assays. Representative images of cells with DAPI nuclear stains in the lower chambers are shown in the *right panels*. Relative invasion ability was calculated using the cell number in the lower chamber of transwell system for each clone compared to that of the vector control. G. Dot plots showing migration velocity of CL1-0 cells measured by transwell migration assays. Relative migration ability was calculated using the cell number in the lower chamber of the transwell system for each sample compared to that of the vector control. H. Diagram of *in vivo* extravasation and lung colonization assay following injection of lung cancer cells into the right ventricle of C57BL/6 mice. Mice were sacrificed 30 mins after intracardial injection. Whole lung perfusion with normal saline was performed to remove blood and cells not adhered to the pulmonary vasculature. I. Numbers of lung cancer cells with FUT4 over-expression (*left panel*, A549_FUT4^high^; *right panel*, CL1-0_FUT4) adhered to the vascular walls or retained in the lung tissues following intracardial injection were visualized under Zeiss Axio Observer microscope. Data information: scale bar in (C, D, F and G); 100 μm. Scale bar in (I); 5 mm. *p* value in (B-D) was calculated by one-way ANOVA with Dunnett’s test, and in (E, F, G, and I) was by Mann-Whitney test. All experiments were performed in three biological replicates and presented as mean ± standard errors. * *p* <0.05, ** *p* < 0.01, *** *p* <0.0005, **** *p* <0.0001. ns: not significant.

Moreover, to confirm whether FUT4-mediated aberrant fucosylation may lead to a stronger cell adhesion—an early step in the metastatic cascade, we performed *in vitro* adhesion assays with several common adhesion molecules within human tissues and blood vessels. Consistent with recent reports^12, 20, 21^, our data showed that FUT4 facilitated binding to E-selectin (on endothelial cells) and L-selectin (on lymphocytes), but not P-selectin (on platelets) or collagen IV (Fig EV1). In light of this, we further performed an *in vivo* assay to investigate the effects of FUT4 on organotropic extravasation ability to model the metastatic process in which the cells travel and lodge themselves at distant sites. We pre-labeled lung cancer cells with CYTO-ID® red tracer dye and injected the cells into the right ventricle of C57BL6 mice. Thirty minutes later, mice were sacrificed and perfused with normal saline intracardially to remove cells not adhered to the pulmonary vasculature (Fig 2H). A marked retention of A549_FUT4^high^ and CL1-0_FUT4 cells in the lungs were observed under a microscope as opposed to their respective vector controls (Fig 2I). These data suggested that FUT4 is a key player in promoting invasion and colonization of lung adenocarcinoma cells.

### FUT4 drives metastasis of lung cancer cells in mouse xenograft models

To substantiate the role of FUT4 on lung cancer metastasis, we investigated whether aberrant FUT4 expression led to spontaneous distant metastasis using two mouse xenograft models. In the tail vein assay for cancer metastasis in nude mice, we observed a rapid lung homing phenomenon enhanced by FUT4. As short as one day post tail vein injection, more A549_FUT4^med^ and A549_FUT4^high^ tumor cells were trapped in the lung areas shown by the IVIS® Spectrum imaging system compared to A549_vector cells (Fig 3A). When we sacrificed mice 28 days post-injections, we found that mice in the FUT4^high^ group had a much greater number of metastatic lung nodules compared to those in the FUT4^med^ and vector groups (p < 0.0005) (Fig 3B). In the second xenograft model, we subcutaneously injected A549_FUT4^med^, A549_FUT4^high^ cells and A549_vector into NOD/SCID immunocompromised mice and evaluated spontaneous metastasis of cancer cells. When mice were sacrificed 56 days post-injection, we observed that mice carrying subcutaneous tumors of A549_FUT4^high^ cells and A549_FUT4^med^ cells had higher frequencies of metastatic lung nodules (Fig 3C and D). Moreover, higher FUT4 levels were correlated with higher metastatic potentials (Fig 3C and D). These data support the significant role of FUT4 in driving metastasis of lung adenocarcinoma cells *in vivo*.

**Figure 3.**
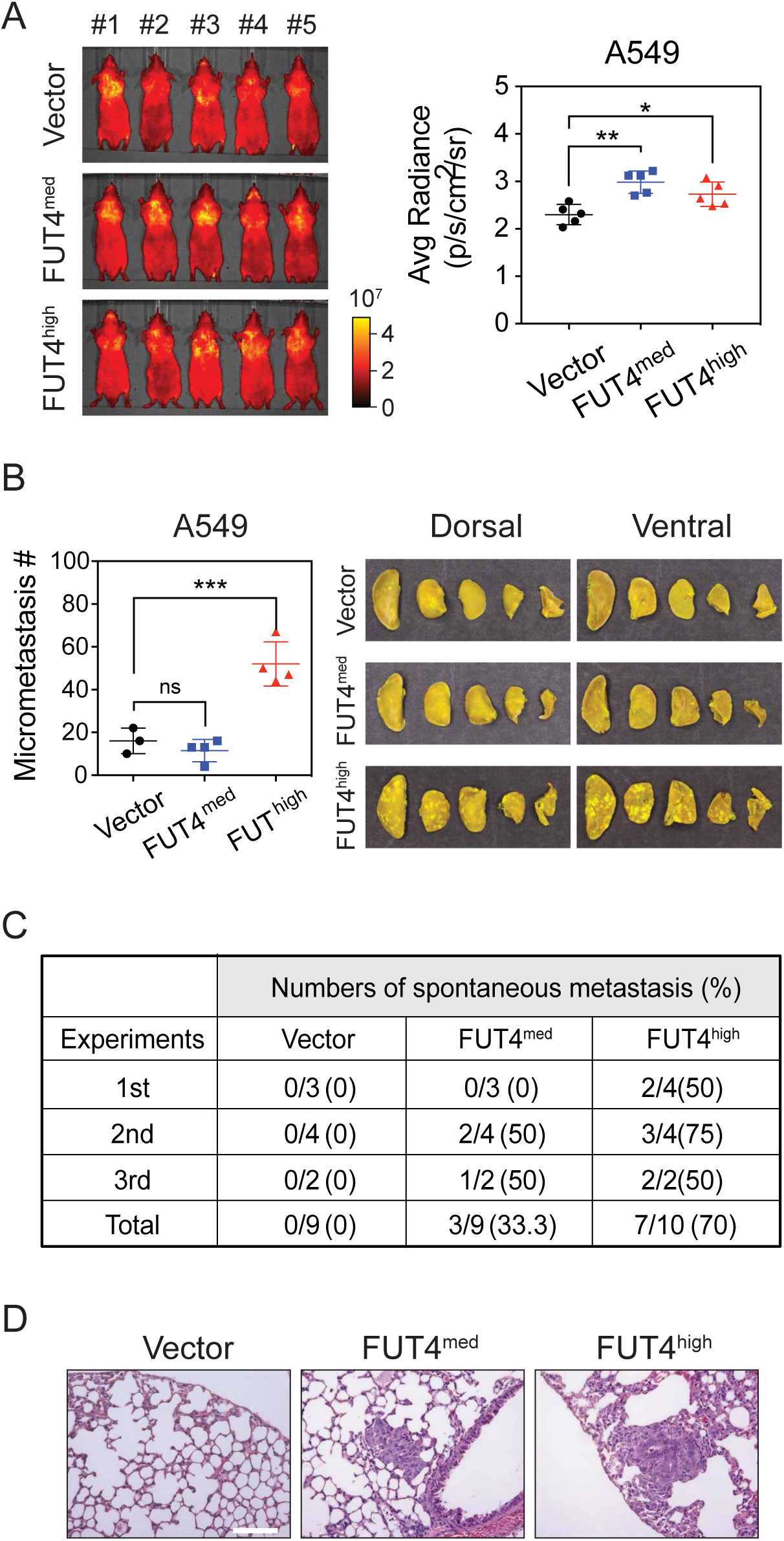
FUT4 potentiates lung homing ability and drives *in vivo* metastasis of lung cancer cells. A. Whole animal imaging of *in vivo* tumor metastasis in nude mice with the IVIS® Spectrum at 24 hours following tail vein injections of A549 lung cancer cells pre-stained with Cyto-ID™ long-term dyes. Quantification of signal radiances of metastatic foci in the dorsal and ventral sides of animals was graphed on the *right panel*. B. Numbers of metastatic foci in the lungs of nude mice at 28 days after tail vein injection of lung cancer cells with FUT4 overexpression (A549_FUT4^med^, A549_FUT4^high^). Pictures of lung nodules from one representative mouse in each group are also shown on the right. C. Numbers (percentages) of mice with spontaneous lung metastases in NOD/SCID mice bearing subcutaneous tumors of A549_vector, A549_FUT4^med^ and A549_FUT4^high^ lung cancer cells. Data are summarized from three independent mouse experiments. D. Representative hematoxylin & eosin (H&E) stained images of lung tissues from mice bearing subcutaneous xenograft lung tumors from (C). Scale bar: 0.1 mm. Data information: *p* value in (A, B) was calculated by one-way ANOVA with Dunnett’s test. * *p* <0.05, ** *p* < 0.01, *** *p* <0.0005. ns: not significant. All experiments were performed in three biological replicates and presented as mean ± standard errors.

### FUT4 mediates activation of membrane trafficking and oncogenic pathways

Fucosylation has been shown to play an important regulatory role in protein structure and functions. An in-depth understanding of global transcriptomic profiling on FUT4-associated cellular processes and pathway alterations may facilitate development of therapeutic strategies to reverse malignant phenotypes. Thus, we performed transcriptomic analysis on A549_FUT4^high^ and CL1-0_FUT4 cells using genome-wide RNA-seq technologies (GEO: GSE120622). We observed that there were 193 and 291 genes significantly upregulated, and 183 and 330 genes significantly downregulated by more than 1.4-fold (p <0.05) between FUT4-overexpressing cells and parental cells in A549 and CL1-0, respectively. For genes commonly regulated in both cell lines, Gene Set Enrichment Analysis^22^ revealed activation of multiple pathways related to membrane trafficking, cell cycle, RNA processing, as well as EGF and TGFβ signaling pathways, among others (Fig 4A and B). The membrane trafficking system mediates intracellular membrane transport of proteins, carbohydrates, lipids between organelles such as endoplasmic reticulum (ER), Golgi complex and plasma membrane^23^. For instance, SEC31A, SEC13, SEC23A, SEC24B, C, D, components of the coat protein complex II (COPII) which facilitates transport vesicle formation from ER, are upregulated in FUT4-overexpressing cells. Similarly, transcriptomic activation of COPA, COPB, COPE, ARCN1, components of another coatomer complex which mediates Golgi outbound cargos, are also noted (Table S2). The data indicate that FUT4-overexpressing cells are in a highly active state of biosynthesis and biomolecule trafficking.

**Figure 4.**
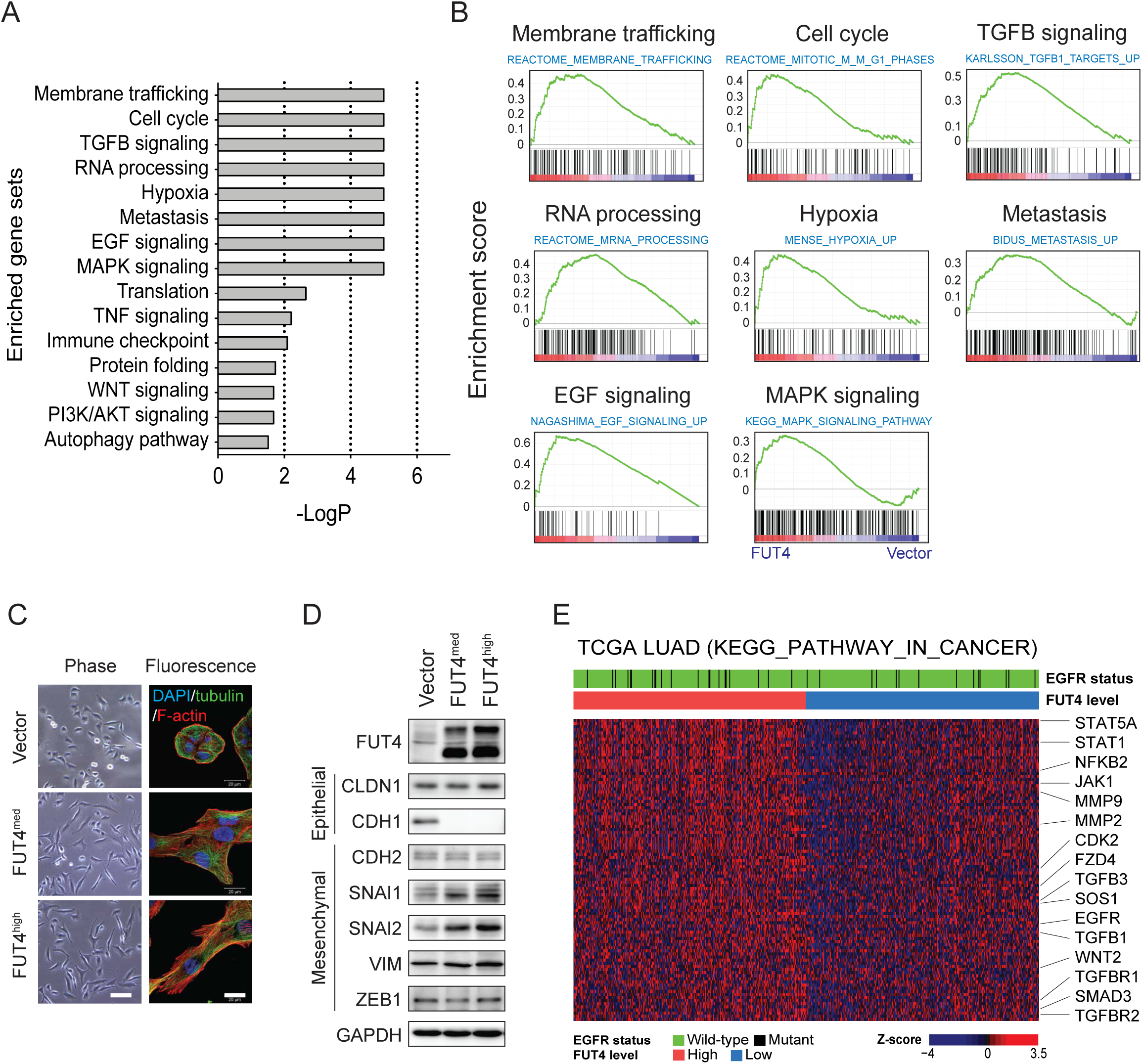
FUT4 mediates activation of membrane trafficking and oncogenic pathways. A. Top 15 positively enriched gene sets in genome-wide RNA-seq data of A549_FUT4^high^ and CL1-0_FUT4 cells versus respective vector controls. Enriched gene set data from Gene Set Enrichment Analysis (GSEA) with nominal *p* value less than 0.05 and false discovery rate (FDR) less than 0.25 are presented. B. GSEA enrichment plots of top 8 positively enriched gene sets in A549_FUT4^high^ and CL1-0_FUT4 cells, including membrane trafficking, cell cycle, TGFβ signaling, RNA processing, hypoxia, metastasis, EGF and MAPK signaling pathways. C. Immunofluorescent imaging analysis of cytoskeleton and cell morphology of A549 lung cancer cells with various levels of FUT4 over-expression (A549_vector, A549_FUT4^med^, and A549_FUT4^high^). Scale bar: 50 μm (*left panel*) and 20 μm (*right panel*). D. Western blot analyses of EMT marker proteins in A549 lung cancer cells with various levels of FUT4 over-expression. E. Heatmap of RNA-seq transcriptomes for the leading edge genes of KEGG_pathway_in_cancer from GSEA analyses of TCGA lung adenocarcinoma. Top sidebars denote mutation status of EGFR and expression levels of FUT4 grouped with the median value cut-off. Data information: *p* value in (A, B) was calculated by one-way ANOVA with Dunnett’s test. * *p* <0.05, ** *p* < 0.01, *** *p* <0.0005. ns: not significant. Experiments were performed in three biological replicates and presented as mean ± standard errors.

In addition, FUT4 appears to modulate pathways related to cell cycle (e.g. CDC6, CDC23, etc), oncogenes (e.g. NRAS, CBFB, FOS), cancer metastasis (e.g. AURKA, CDK1, PLK4), as well as major oncogenic signaling pathways – TGFβ, EGF, MAPK, and WNT signaling (Fig 4A, B, and Table S2). As a functional validation to activating TGFβ and EGF signaling in FUT4-overexpressing cells, we demonstrated that FUT4 leads to morphological changes characteristic of EMT, a phenomenon mediated by the two signaling pathways, with a prominent reorganization of the cytoskeleton toward a more mesenchymal phenotype in A549_FUT4^med^, A549_FUT4^high^ and CL1-0_FUT4 cells (Fig 4C and EV2). Consistently, these changes are associated with down-regulation of CDH1 (E-cadherin) and up-regulation of SNAI1 (a.k.a SNAIL) and SNAI2 (a.k.a SLUG), two EMT-inducing transcription factors (Fig 4D). Interestingly, minimal increases of mesenchymal proteins—VIM (vimentin) and CDH2 (N-cadherin) are noted, suggesting FUT4 induces an intermediate EMT phenotype in A549 lung cancer cells (Fig 4D).

To investigate whether these FUT4-mediated pathway alterations were also shown in primary tumor tissues, we examined transcriptomic data in the TCGA cohort of 517 lung adenocarcinomas. In consistent with cell line data, we found that high FUT4-expressing tumors displayed significantly enhanced activities in oncogenic signaling networks including TGFβ, EGF, MAPK, and WNT as opposed to low FUT4-expressing tumors (Fig 4E). Collectively, our analysis suggests that FUT4 induces global reprogramming of the transcriptomic program towards a more malignant phenotype via concurrent alterations of multiple cellular processes and signaling pathways

### FUT4 induces aberrant fucosylation of intracellular transport and signaling proteins

Next, we investigated how FUT4 and its fucosylated glycans modulate activities of global cellular processes and signaling pathways. Among the three Lewis antigens whose synthesis may involve FUT4 — Lewis x (Le^x^), Lewis y (Le^y^) and sialyl-Lewis x (sLe^x^)^4, 24–26^, we identified Lewis x (Le^x^) as the major glycan antigen synthesized by FUT4 using flow cytometric analysis, which showed that Le^X^-expressing cell populations were markedly increased in A549_FUT4^med^, A549_FUT4^high^ cells and CL1-0 cells, whereas Le^y^ and sLe^x^ were not altered (Fig 5A and EV3). Moreover, immunoblotting with anti-Le^x^ showed global increases of Le^X^-bearing glycoproteins in the whole cell lysate of FUT4-overexpressing A549 and CL1-0 cells (Fig EV3).

**Figure 5.**
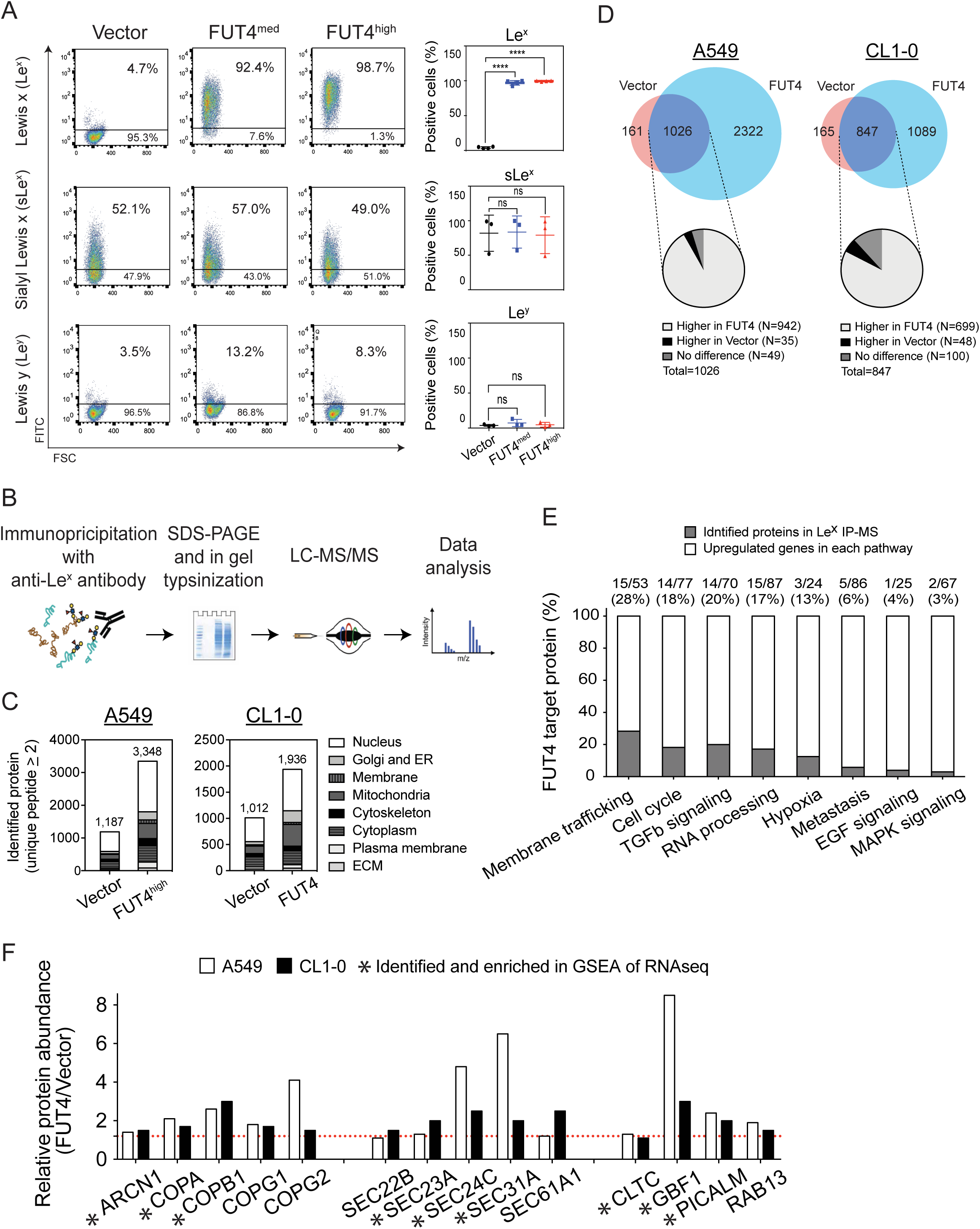
FUT4 induces aberrant fucosylation of intracellular transport and signaling proteins. A. Flow cytometric analysis of cell surface glycans, Lewis x (Le^x^), sialyl Lewis x (sLe^x^) and Lewis y (Le^y^) in FUT4-overexpressing A549 lung cancer cells. *Left panels*, representative flow cytometric dot plots of Le^x^, sLe^x^ and Le^y^ expression are shown. *Right panels*, dot plots showing percentages of cells expressing individual surface glycans in three biological replicates. B. Experimental diagram of immunoprecipitation (IP) with anti-Le^x^ antibody followed by liquid chromatography-mass spectrometry (LC/MS-MS). C. Subcellular localizations of FUT4 acceptor proteins bearing higher levels of Le^x^ antigens in A549_FUT4^high^ and CL1-0_FUT4 cells compared to their respective vector controls. Y axis denotes numbers of identified protein in each subcellular location. D. Venn diagram comparing the numbers of identified proteins in FUT4-overexpressing cells vs vector controls in A549 (*left panel*) and CL1-0 (*right panel*) cells. Pie charts show percentages of increased or decreased fucosylated proteins in FUT4-overexpressing cells. E. Bar graph reveals the percentages of FUT4 acceptor proteins (gray bars) in the core enriched genes of top 8 positively enriched gene sets upon FUT4 overexpression. The numbers above the bars denote the numbers (numerator) of identified proteins by anti-Le^x^ IP-MS and the numbers (denominator) of core enriched genes in individual gene sets. F. Bar graphs show relative abundance of membrane trafficking-related proteins in A549_FUT4^high^ and CL1-0_FUT4 cells compared to their respective vector controls. The proteins also identified in RNA-seq GSEA analysis are marked with asterisks below.

To delineate the cellular effects of FUT4-mediated fucosylation, we went on to probe acceptor proteins fucosylated by FUT4 using immunoprecipitation (IP) with anti-antibody followed by tandem mass spectrometry (LC-MS/MS)(Fig 5B). We identified 3,348 and 1,936 proteins bearing Le^x^ antigen in A549_FUT4^high^ and CL1-0_FUT4 cells, and 1,187 and 1,012 proteins in their respective vector controls (Fig 5C) (MassIVE: MSV000083028). We found that FUT4 synthesized Le^x^ on a wide variety of proteins in different intracellular compartments (Fig 5C). Among the Le^x^-bearing proteins upregulated in A549_FUT4^high^ and CL1-0_FUT4 cells (Fig 5D), many of them participate in the activated cellular processes and signaling pathways revealed by RNA-seq, including membrane trafficking, cell cycle, TGFβ signaling, etc (Fig 5E). The data suggest a strong link between FUT4-mediated aberrant fucosylation and heightened activities of cellular processes and signaling. In particular, 28% of upregulated genes that drive the enrichment of the membrane trafficking process are aberrantly fucosylated in FUT4-overexpressing cells (Fig 5E). In particular, components of a coatomer complex—ARCN1, COPA1, COPB1, as well as SEC family proteins that form the coat protein complex II (COPII)—SEC23, 24, 31A, 61A1, are highly fucosylated (Fig 5F). Moreover, aberrant fucosylation of RAB13, a small GTPase regulating vesicular trafficking between trans-Golgi network and recycling endosomes^27^, is also noted (Fig 5F). These data strongly suggest that FUT4 high-expressing cells have a disturbed intracellular transport state.

### FUT4 enhances metastasis-related signaling via fucosylation of cascade proteins

In addition to facilitating intracellular transport, FUT4-fucosylated proteins are simultaneously involved in several major signaling pathways known for metastasis, angiogenesis and EMT, including EGF, TGFβ, WNT and HIPPO pathways (Fig 6A). Remarkably, these FUT4 acceptor proteins interact physically and form an intricate network of key mediator proteins at multiple levels downstream of these pathways (Fig 6B). To further elucidate the potentiating effect of FUT4-mediated aberrant fucosylation on signal transduction, we found that FUT4 high-expressing cells exhibit enhanced signaling activity as demonstrated by an increase of phospho-ERK (pERK) and phosphor-Smad2 (pSmad2) at 0.5 to 2 hours post EGF (Fig 6C and D) or TGFβ (Fig 6E and F) pathway activation in A549_FUT4^high^ compared to the vector control (Fig 6C and D). Similar phenomenon was also observed in CL1-0_FUT4 cells (Fig EV4). This suggest that FUT4-mediated cascade protein fucosylation augments signal transduction upon pathway activation.

**Figure 6.**
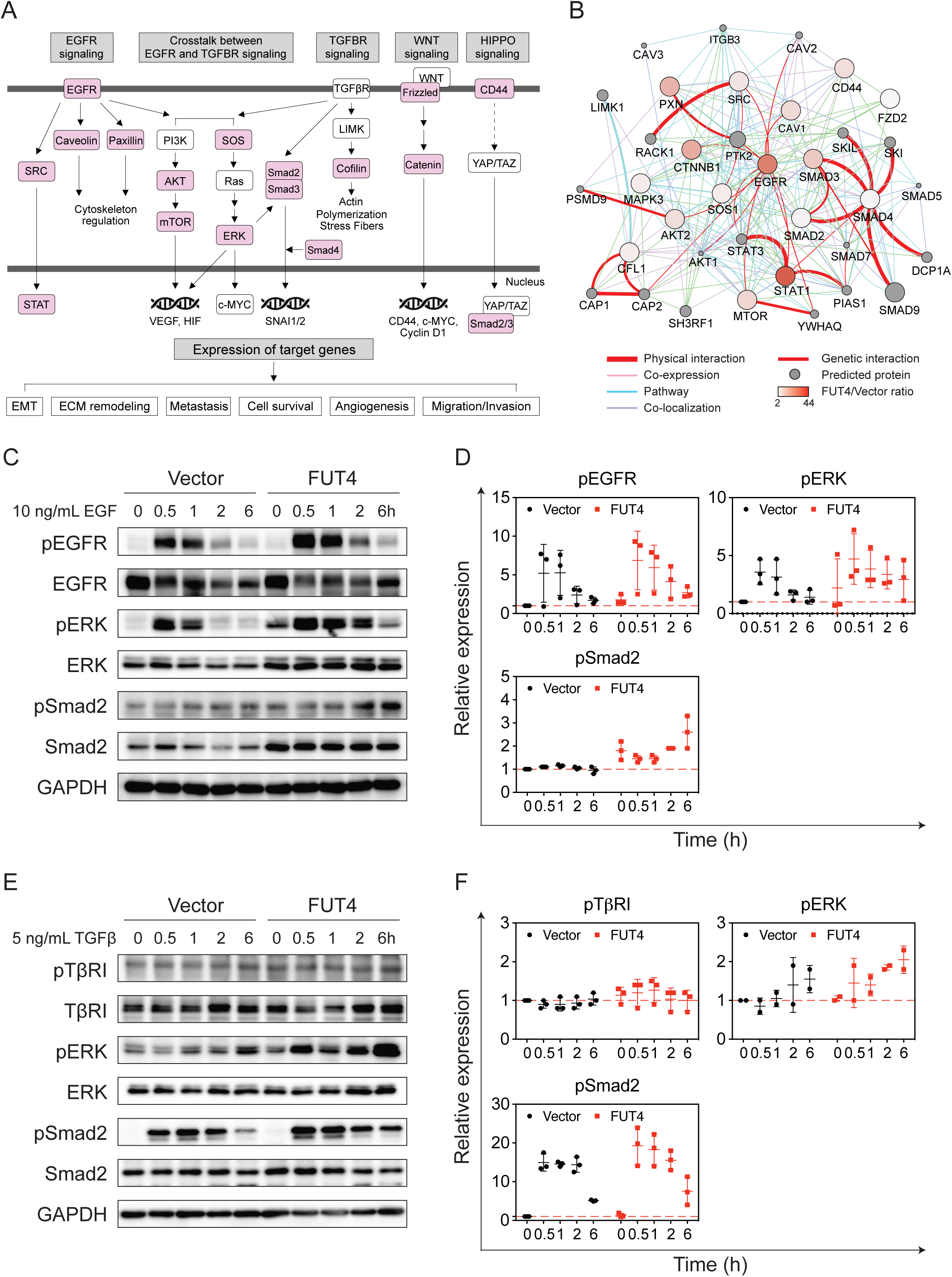
FUT4 enhances metastasis-related signaling via fucosylation of cascade proteins. A. Immunoprecipitation (IP) with anti-Le^x^ antibody followed by tandem mass spectrometry (LC-MS/MS) in A549_FUT4^high^ and CL1-0_FUT4 cells reveals key mediator proteins (in pink) in oncogenic signaling cascades including EGF, TGFβ, WNT and HIPPO pathways. B. Protein networks showing protein-protein interactions between FUT4 acceptor proteins bearing higher levels of Le^x^ antigens in A549_FUT4^high^ cells relative to A549_vector cells. The data were analyzed by Cytoscape (ver. 3.6.1) using GeneMania application (ver. 3.4.1). C. Western blot analyses of signaling cascade proteins in the EGFR pathway in A549_FUT4^high^ versus A549_vector cells at 0, 0.5, 1, 2, and 6 hrs following the addition of 10 ng/mL EGF. Quantifications of signal intensities on western blots for phospho-EGFR, phospho-ERK and phospho-Smad2 from three biological replicates are graphed on the *right panel*. D. Western blot analyses of signaling cascade proteins in the TGFβ signaling pathway in A549_FUT4^high^ versus A549_vector cells at 0, 0.5, 1, 2, and 6 hrs following the addition of 5 ng/mL TGFβ. Quantifications of signal intensities on western blots for phospho-TGFβ receptor I, phospho-ERK and phospho-Smad2 from three biological replicates are graphed on the *right panel*. Data information: *p* value in (D, F) was calculated by one-way ANOVA with Dunnett’s test. * *p* <0.05, ** *p* < 0.01, *** *p* <0.0005. ns: not significant. All experiments were performed in three biological replicates and presented as mean ± standard errors.

### Genetic depletion of FUT4 diminishes *in vitro* and *in vivo* aggressive phenotypes of human lung cancer cells

To investigate the potentials of FUT4 as a therapeutic target, we knocked down FUT4 using an shRNA approach in CL1-5 lung adenocarcinoma cells, which have aggressive migration/invasion behaviors characterized in the previous study^28^, and a relatively high expression of FUT4 (Fig 2A and 7A). Migration and invasion abilities of cancer cells appeared to be reduced in CL1-5_shFUT4#751 cells which contained the lowest expression level of FUT4 among all shRNA clones (#751, #753 and #792) compared to the vector control (Fig 7B and 7C). The other two knockdown clones (#753 and #792) also showed a consistent statistical trend towards significance (Fig 7B and 7C). Moreover, knock-down of FUT4 significantly decreased the extravasation and retention ability of CL1-5 cells in the pulmonary vasculature when we injected the cells into the right ventricle of C57BL6 mice to evaluate their organotropic extravasation ability (Fig 7D). Furthermore, when we silenced FUT4 using shRNA in CL1-5 lung cancer cells which have spontaneous *in vivo* metastatic ability^28^, we found that FUT4 silencing markedly abolished metastasis of lung cancer cells (Fig 7E).

**Figure 7.**
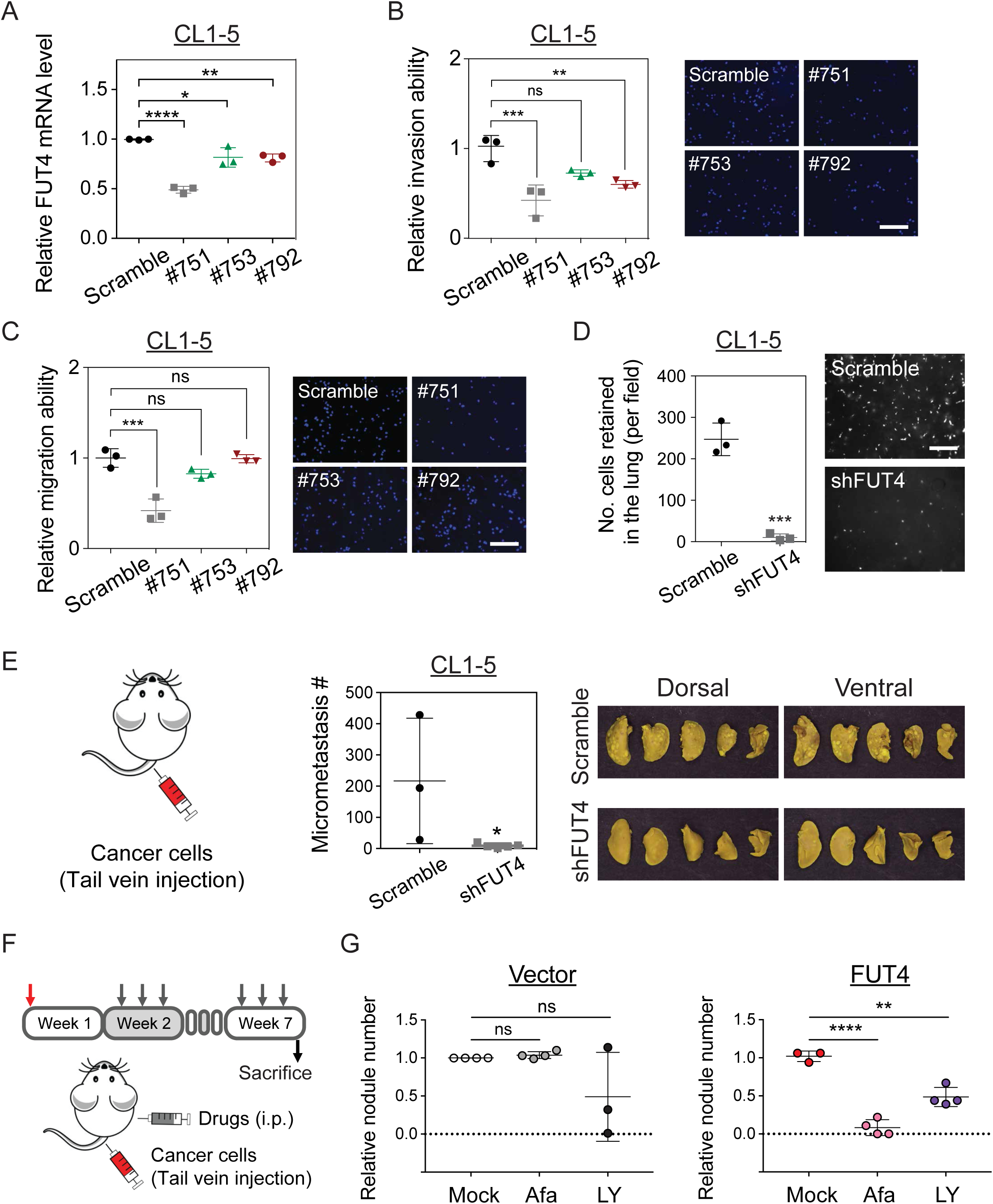
Genetic depletion of FUT4 diminishes *in vitro* and *in vivo* aggressive phenotypes of human lung cancer cells. A. FUT4 mRNA levels in CL1-5 lung cancer cells transfected with FUT4 shRNA (#751, #753, #792) measured by quantitative real-time PCR. B. Dot plots showing relative invasion ability of CL1-5 cells with FUT4 knockdowns measured by matrigel-based transwell invasion assays. Representative images of cells with DAPI nuclear stains in the lower chambers are shown. C. Dot plots showing relative migration ability of CL1-5 cells with FUT4 knockdowns. Relative migration ability was calculated using the cell number in the lower chamber of the transwell system for each clone compared to that of the scramble control. Representative images of cells with DAPI nuclear stains in the lower chambers are shown. D. Numbers of lung cancer cells with FUT4 knockdowns (CL1-5_shFUT4) adhered to the vascular walls or retained in the lung tissues following intracardial injection visualized under Zeiss Axio Observer microscope. Scale bar: 5 mm. *p* value was calculated by Mann-Whitney test. *** *p* <0.0005. E. Numbers of metastatic foci in the lungs of nude mice at 28 days after tail vein injection of lung cancer cells with FUT4 knockdown (CL1-5_shFUT4). Pictures of lung nodules from one representative mouse in each group are shown in the *right panel*. F. Diagram of *in vivo* lung cancer metastasis assay via tail vein injection in nude mice subject to treatments with EGFR or TGF-β inhibitors. G. Dot plots showing relative metastatic abilities of A549_vector or A549_FUT4^high^ lung cancer cells in nude mice receiving 0.25 mg/kg Afatinib or 15 mg/kg LY2157299 treatment. Numbers of metastatic foci in the lungs were counted and normalized to the numbers in the mock (normal saline)-treated groups. Data information: *p* value in (A, B, C, G) was calculated by one-way ANOVA with Dunnett’s test. * *p* <0.05, ** *p* < 0.01, *** *p* <0.0005, and in (D, E) was calculated by Mann-Whitney test. ns: not significant. Data in (A-F) were performed in three biological replicates and presented as mean ± standard errors.

In addition, to test whether FUT4-mediated malignant phenotype could be pharmacologically diminished by targeting signaling networks provoked by FUT4, we performed *in vivo* lung cancer metastasis assay via tail vein injection of A549_vector or A549_FUT4^high^ cells into nude mice, which subsequently received intraperitoneal treatment of either 0.25 mg/kg afatinib (an EGFR inhibitor) or 15 mg/kg LY2157299 (a TGF-β inhibitor) three times a week (Fig 7F). We found that both drugs partially diminished lung metastasis in A549_FUT4^high^ cells (Fig 7G). Interestingly, afatinib demonstrated differential inhibitory effects between A549_FUT4^high^ and A549_vector cells (Fig 7G), which implies A459_FUT4^high^ cells have higher dependency on EGF signaling for distant metastasis. These proof of principle data indicate that targeting FUT4-potentiated pathways may provide a therapeutic opportunity against metastasis of FUT4-high-expressing tumors, and that combination therapy against multiple FUT4-medicated cellular processes may be required to achieve better efficacy in patients with FUT4-high-expressing tumors.

## Discussion

Aberrant expression of fucosylated epitopes such as Lewis antigens have been widely reported in various types of cancer^4^. These Lewis antigens are mainly synthesized by α1-3/ α1-4 fucosyltransferases in a tissue-specific manner. In the current study, we holistically evaluated all α1-3/ α1-4 fucosyltransferases and identified FUT4 as a prognostic biomarker in Asian and Western cohorts of lung cancer patients through large scale transcriptomic analysis. Asian lung cancer is known to be a molecularly and etiologically distinct identity composed of a much higher percentage of never smokers and EGFR mutations^17, 18^ as compared to the western counterpart^15, 16^. Yet, comparative transcriptomic analyses of Eastern and Western cohorts revealed striking similarities in the expression patterns of α1-3 terminal fucosyltransferases in non-small cell lung cancers despite differences in the mutational status and ethnicities. Among all α1-3 FUTs, FUT3, 4, 6, 10 and 11 appeared to be highly expressed in both lung adenocarcinoma and squamous cell carcinoma. Moreover, despite the general conception that FUTs may facilitate cell proliferation or invasion in cancers^19, 29–31^, our genome-wide analysis in large patient cohorts revealed that FUTs may also have protective prognostic effects, which is generally underappreciated. In our data, while FUT4 is correlated with poor survival, FUT1 and FUT10 appeared to confer survival benefit in lung cancer. It is interesting that FUT1 has been shown to promote proliferation and spreading potential of ovarian carcinoma^32, 33^. This suggests that the same fucosyltransferase may act on different sets of acceptor proteins in different tumor types. While we delineated the role of FUT4 in lung adenocarcinoma, detailed mechanistic and clinical studies may be required to elucidate the precise roles of other FUTs in individual tumor types.

Furthermore, our combined transcriptomic and glycoproteomic approach provides a comprehensive view of cellular processes and intracellular signaling altered by FUT4. This glycosylation-related reprograming of cellular transcriptome collectively contribute to the malignant phenotype of cancer cells, instead of one or two pathways being held accountable as shown in previous reports^11, 19, 34, 35^. In addition to major oncogenic pathways such as EGF/MAPK, NF-κB/p65, and TGF-beta pathways, our data reveal that FUT4 induces an active intracellular trafficking state as the top enriched cellular process. Notably, the alterations of intracellular vesicle trafficking have recently been shown to drive oncogenesis and regulate cancer behavior^36, 37^. FUT4 not only upregulates but also aberrantly fucosylates many components of vesicle transport proteins including SEC protein family in COPII complex, and regulators of vesicle trafficking such as small GTPases (Fig 5F). The hyperactive intracellular transport system may underly the neoplastic phenotypes including proliferation, migration and invasion^36, 37^. Moreover, it is increasingly acknowledged that intracellular transport and signaling pathways (e.g. PI3K-Akt, EFGR and FGFR signaling) are interconnected^38^. COPII trafficking was shown to involve in regulating surface levels of receptor tyrosine kinases^39^. In our data, overexpression of FUT4 markedly reinforces the crosstalks between membrane trafficking and signal transduction. This provides a molecular basis for targeting these pathways as novel therapeutic strategies in treating high FUT4-expressing tumors.

In addition, while fucosylation of surface receptors, including EGFR, has been reported^6, 14^, attempts to map intracellular targets of FUTs for a deeper mechanistic interrogation are fairly limited due to transient interactions between FUTs and their acceptor proteins as they fold and pass through the Golgi apparatus. A global view on the complex protein network crosstalks has been lacking. Here, we took a glimpse into candidate FUT4 acceptor proteins through proteomics approach by tandem mass spectrometry (LC-MS/MS) based on the presence of Le^x^, a predominant product synthesized by FUT4 in lung cancer cells in our system (Fig 5). We discovered an intricate network of Le^x^-decorated intracellular proteins involved in membrane trafficking between endoplasmic reticulum, trans Golgi networks, and endosomes, etc. These membrane trafficking network and major signaling pathways are closely interconnected and co-activated in FUT4 over-expressing cells on both transcriptomic and protein levels. Notably, we demonstrated that fucosylated cascade proteins in signaling pathways such as EGF or TGFβ lead to enhanced signal transduction upon pathway activation with exogenous ligands. Our finding brings to light the additional biological role of Le^x^ or other carbohydrate epitopes on cytosolic proteins aside from being merely a surface antigen, which may open up research on whether and how altered fucosylated glycans modulate the activities of intracellular proteins. Further studies such as glycosylation/fucosylation site-directed mutagenesis are warranted to decode how Le^x^ can alter protein functions and stability.

Most importantly, our *in vitro* and *in vivo* studies demonstrated a prime role of FUT4 during the process of lung cancer metastasis, which highlights the potential of FUT4 as an attractive therapeutic target. Indeed, knockdown of FUT4 in our study significantly diminished lung metastases in mouse xenograft models without apparent toxicities. Currently, glycosylation-related therapeutics is under active development, ranging from modulating stability of therapeutic proteins via glycosylation to glycoprotein-based biodrugs^40^. Multiple attempts have been taken to identify inhibitors of sialyl- and fucosyl-transferases via high-throughput screening^41–43^. Non-selective metabolic inhibitors such as synthetically-derived fucose-related molecules have been shown to inhibit global fucosylation thereby diminishing neutrophil extravasation or xenograft tumor growth in immunocompromised mice^44^. As our data showed that individual fucosyltransferases may have opposite effects on patient prognosis, we think that development of FUT4-specific inhibitors may be necessary for preventing lung cancer metastases, and at the same time minimize potential off-target effects due to non-selective ablation of global fucosylation.

On the other hand, for a broader clinical implication, we show that FUT4-mediated EGFR or TGFβ activation can occur in lung tumors not carrying pathway-specific driver mutations. As shown in the current study, EGFR- or TGFβ-targeted therapy had therapeutic efficacy to a certain extent in FUT4 high-expressing cells. This indicates that pharmacologically targeting FUT4-mediated pathways, such as membrane trafficking, EGF, TGFβ, WNT signaling, may be a reasonable therapeutic option alternative to FUT4 depletion therapy for treating a subset of EGFR-wild type lung cancers with medium-to-high FUT4 expressions. As inhibitors targeting these FUT4-mediated oncogenic signaling pathways are either ready for trials or FDA-approved for clinical use, it would be rather straightforward to design future clinical trials and test the use of these targeted therapies in patients with wild-type EGFR lung cancers expressing higher levels of FUT4.

In summary, our results suggest that FUT4, a terminal α1,3-fucosyltransferase, is an important prognostic indicator in lung adenocarcinoma patients. FUT4 promotes lung cancer progression at multiple steps of metastatic process — increased migration/invasion, vascular adhesion, extravasation, and tumor metastasis at distant organs. These profound cellular and functional effects can be attributable to the augmentation of global transcriptomic alterations such as intracellular membrane trafficking, oncogenic signaling pathways, which may be reversed by targeting FUT4 directly or pathway-specific inhibitors. Our study may pave way to development of novel glycol-based therapeutic strategies for clinical management in lung cancer metastases.

## Materials and methods

### TCGA data acquisition and analysis

Normalized RNA-seq expression datasets and EGFR mutation status of The Cancer Genome Atlas (TCGA) lung adenocarcinoma (LUAD) and lung squamous cell carcinoma (LUSC) cohorts were acquired from Firehose data repository using the R/Bioconductor package “RTCGAToolbox” (version 2.10.0, run data 2016-01-28)^45^. We used log-rank test to compare between-group survival differences. Patients were censored at the last follow-up or death, with post-surgery recurrence (>3 months) classified as an event for relapse-free survival. As for overall survival, patients were censored at the last follow-up, and death was classified as an event. Heatmaps of gene expression profiles were created by R package “pheatmap” version 1.0.10.

### Human tissues and subject follow-up data

Surgically-resected lung tumors and adjacent uninvolved lung tissues were obtained from patients with non-small cell lung cancer from National Taiwan University Hospital (NTUH, Taipei, Taiwan) from 1994 to 2010. Clinical information including gender, histological type, stage, and overall survival was collected. Median follow-up time was 51 months (range 2-164 months). The staging of lung cancer was based on the American Joint Committee on Cancer (AJCC) TNM system. This study has been approved by the Institutional Review Board (IRB) of National Taiwan University Hospital (Approval No.: 201701010RINB).

### Quantitative PCR analysis

PCR was carried out on a CFX96 TouchTM Real-Time PCR Detection System using Sybr Green I detection (BIO-RAD) according to the manufacturer’s instructions. Relative expression levels were normalized to TBP (TATA-box-binding protein) and calculated using the 2^−ΔCt^ method as described. Primers were purchased from IDT (Integration DNA Technologies) and the sequences are listed in Table S4. mRNA expression levels of FUT4 were measured in 107 human lung adenocarcinoma tissues obtained from NTUH and matched with their clinical data.

### Human lung cancer cell lines

CL1-0 and CL1-5 human lung adenocarcinoma cell lines were kindly provided by Prof. Pan-Chyr Yang (Department of Internal Medicine, National Taiwan University Hospital, Taipei, Taiwan). The CL-1 cell line was established from a 64-year-old man with a poorly differentiated adenocarcinoma^28^. The CL1-5, a subclone derived from CL1-0, possesses higher invasiveness and metastatic ability *in vitro* and *in vivo*. A549 human lung adenocarcinoma cell line was obtained from ATCC. Cell line authentication was performed by STR analysis for all cell lines used in this study. Cells were maintained in complete RPMI-1640 media with 10% (v/v) FBS, 1% (v/v) L-glutamine, and 1% (v/v) Penicillin-Streptomycin (Gibco™) at 37°C and 5% CO_2._ Detection of mycoplasma in cell culture was performed every season in our lab to avoid mycoplasma contamination. For signaling transduction assays, cells were cultured in medium containing 0.5% (v/v) FBS for 16 hours for cell cycle synchronization. Then cells were treated with EGF (10 ng/mL; AF-100-15, PeproTech) or TGF-β1 (5 ng/mL; 100-21, PeproTech) for 1, 2, 6 hours, respectively. To block the effects of growth factors, cells were pre-treated with Afatinib (2 μM; T2303, TargetMol) or LY2157299 (10 μM; T2510, TargetMol) for 1 hour, followed by treatment with EGF or TGF-β for 24 hours.

### RNA interference

FUT4 expression was knockdown using shERWOOD UltramiR lentiviral inducible shFUT4 system. pZIP-Mcmv-shFUT4 plasmids (15 μg) were transfected into cells (5*106 cells) using Lipofectamine™ 3000 transfection reagents (45 μL). Cells were incubated for 16 hours and selected by fresh RPMI medium with 2 μg/mL puromycin for 24 hours. The effects of shRNA knockdown will be evaluated using proliferation assays, migration assays, and invasion assays.

### Single cell tracking

Cells were seeded on the 6-well plate overnight. On the next day, time-lapse imaging of the cells was performed using an automated inverted microscope (Leica DMI 6000B) for 24 hours. Data were analyzed by MetaMorph^®^ software. For each sample, at least 20 cells were selected and tracked for cell motility calculation.

### Invasion/migration assay

Cell invasion assays were performed using 24-well transwell units with permeable polycarbonate filter with pore sizes of 8 μm. 24-well transwell units were pre-coated with 2 mg/mL BD Matrigel^®^ (BD Biosciences), and incubated at 37oC for 1 hour. Each lower compartment of the transwell unit contained 10% FBS as chemoattractant. 5*10^4^ Cells in 0.5 ml RPMI with 0.5% BSA were added into the upper compartment of the transwell unit and were incubated at 37oC overnight. On the next day, the Matrigel^®^ coated on the filter was removed with cotton swabs, and the filter was gently separated from the transwell unit. The cells attached to the lower surface of the filter membrane were fixed with 4% paraformaldehyde in PBS, and stained with 300nM DAPI (4’,6-Diamidino-2-Phenylindole, Dihydrochloride). The number of cells was counted at 400X magnification under a fluorescent microscope. For each sample, the average number of cells from eight high-power fileds (HPF) was recorded. All experiments were performed in triplicate. Cell migration assays were performed using the same 24-well transwell units without the Matrigel^®^ coating.

### Mouse experiments

4-6 weeks old male C57BL/6JNarl, NOD.CB17-*Prkdc^scid^*/JNarl (NOD SCID) and BALB/cAnN.Cg-*Foxn1nu*/CrlNarl (NUDE) mice were purchased from National Laboratory Animal Center (Taiwan) and maintained under standard pathogen free conditions. *<u>In vivo adhesion/extravasation assays:</u>* A549 (FUT4^high^ and vector control) and CL1-5 (shFUT4 and scramble control) human lung cancer cells were pre-stained with Cyto-ID^™^ long-term dyes (Enzo Life Sciences), and intracardially injected into the right ventricle of 6-week-old C57BL/6JNarl mice (5 mice per group; 5*10^5^ cells/mouse). Mice were kept warm with electric heating blankets for 20 minutes. Mice were then sacrificed and intravenously perfused with phosphate-buffered saline to remove blood cells and unattached human cancer cells. The lungs were removed and fixed with 1% agarose on a 35mm imaging μ-Dish with a high glass bottom (ibidi, 81158). The lungs were examined by the Axio Observer Inverted Microscope System (Zeiss). *<u>Tail Vein Assay for Cancer Metastases:</u>* A549 (FUT4^high^ and vector control) and CL1-5 (shFUT4 and scramble control) human lung cancer cells were pre-stained with Cyto-ID^TM^ long-term dyes (Enzo Life Sciences), and were injected into the tail vein of 6-week-old nude mice with 5*10^5^ cells per mouse. In vivo bioluminescent imaging to monitor cancer cell metastasis was performed using In Vivo Imaging System (IVIS) at day 1 and day 7 after tail vein injection. Mice were euthanized with carbon dioxide at 28 days. All organs were removed and fixed in 10% formalin. The lung nodules were counted through gross inspection and under microscopic examination. The number of mice used in each group (n = 6) was based on the goal of having 98% power to detect a 2-fold difference in nodule numbers between groups at p < 0.05. *<u>Spontaneous metastasis of subcutaneous tumor xenografts:</u>* Tumor xenografts were established in 6-week-old NOD SCID mice via subcutaneous injection into the dorsal region of each mouse with 1 x 10^7^ cells/0.15 ml Hank’s balanced salt solution containing Matrigel^®^ (BD Biosciences). Experiments were performed at five mice in each group (vector control, FUT4^med^ and FUT4^high^). Mice were then sacrificed at 8 weeks. Distant metastases in the lungs were evaluated by gross visualization of nodules on the lung surfaces, and by microscopic imaging of lung tissue sections. *<u>Drug treatment assay:</u>* A549 (FUT4^high^ and vector control) human lung cancer cells were injected into the tail vein of 6-week-old nude mice with 5*10^5^ cells per mouse to establish lung cancer metastatic model. Mice were intraperitoneally injected with DMSO (mock control), afatinib (0.25 mg/Kg BW) (TargetMol TM-T2510) and LY2157299 (15 mg/Kg BW) (TargetMol TM-T2303) three times per week from week 2 to week 7 post-injection to treat metastatic lung tumor. Mice were euthanized with carbon dioxide at 49 days. Lung were removed and fixed in 10% formalin. The lung nodules were counted through gross inspection and under microscopic examination. The number of mice used in each group (n = 6) was based on the goal of having 98% power to detect a 2-fold difference in nodule numbers between groups at p < 0.05.

### Adhesion assay

The uncharged polystyrene 96-well plate were coated with different recombinant proteins-Collagen IV (Merck Millipore), P-Selectin, E-Selectin or L-selectin (PeproTech), and incubated at room temperature for 2 hours. The coated 96-well plates were blocked with 5% BSA/PBS at 37oC for 1 hour. Subsequently, 5 x 10^4^ cells in serum-free medium were added onto the coated 96-well plate. After incubation at 37oC for 1 hour, non-adherent cells were gently washed away twice with 1x PBS. Adherent cells were fixed with 4% paraformaldehyde/PBS for 20 minutes, and stained with 300 nM DAPI. Cell images were obtained with MD ImageXpress Micro XL High-Content Analysis System (Molecular Devices). Numbers of adherent cell were calculated with the MetaMorph software (Molecular Devices).

### RNA-sequencing and data analysis

Total RNAs were isolated from human lung cancer tissues from National Taiwan University Hospital and lung cancer cell lines. RNA integrity was evaluated with an Agilent 2100 Bioanalyzer, and were subjected to quality check with a Bioanalyzer 2100 using an RNA 6000 nano kit (Agilent Technologies). RNA sequencing was performed using Illumina HiSeq 4000 ® (for lung cancer tissue RNAs) and Illumina NextSeq 500 ® systems (for cell line RNAs). Raw reads in FASTAQ format were processed using Trimmomatic software version 0.33 for adapter trimming and quality filtering ^46^. The processed sequencing reads were mapped to the human reference genome (hg19) using bowtie v2.2.6 with parameters: --forward-prob 0 --output-genome-bam for cell lines and clinical samples ^47, 48^. The mapped reads in BAM files were annotated with GENCODE Release 25 (GRCh37), using GenomicFeatures (version 1.32.2) and GenomicAlignments (version 1.16.0) packages. Finally, DESeq2 package (version 1.20.0) was used to generate a raw-counts matrix followed by FPKM normalization and differential expression analysis ^49^. Gene Set Enrichment Analysis (GSEA) was performed using the official software javaGSEA version 3.0 with the Molecular Signatures Database (MSigDB, version 6.1) containing BioCarta, Reactome and KEGG gene sets ^22, 50^. The *signal-to-noise metric* to rank differentially-expressed genes in GSEA. Number of permutations were set at 1000, and enrichment statistic were set as weighted. The resulting gene sets with nominal *p* value < 0.05 were considered as enriched. All RNA-seq data have been deposited in GEO: GSE120622.

### Immunofluorescence staining

For immunolocalization, samples were blocked with 1% (wt/vol) BSA in PBS for 1 hour, followed by incubation at 4°C with primary antibodies for 16 h and with secondary antibodies (donkey anti-mouse IgG Alexa 488 or donkey anti-goat IgG Alexa 488, 1:250 dilution) for 1 hour. Cells were counterstained with 300 nM DAPI for 10 min before mounted with ProLong Gold Antifade Mountant (ThermoFisher Scientific). Images were obtained with the Zeiss LSM 880 confocal microscope, and analyzed with ZEN microscope software. Antibodies used in this study were listed in Supplemental Table 2.

### Immunoprecipitation

Lewis and TβRI immunoprecipitations were performed using PierceTM Protein L Magnetic Beads (ThermoFisher Scientific, #88849) and PierceTM Protein A/G Magnetic Beads (ThermoFisher Scientific, #88802) respectively, according to the manufacturer’s protocol. A549 and CL1-0 cells (both vector control and FUT4-overexpressed cells) were lysed with RIPA buffer containing a protease inhibitor cocktail (GOAL Bio), and incubated on ice for 20 minutes. Cell lysate was centrifuged for 15 min at 4°C, and an aliquot of the supernatant was kept aside on ice as input. 1 mg of cell lysate was incubated with 40 μg of primary antibody per IP at 4oC on a shaker overnight. The following antibodies were used for IP: TGF beta receptor I (Abcam, ab31013) and Lewis X (Abcam, ab3358). Next day, the magnetic beads were loaded in the cell lysate for 4 hours at room temperature and washed according to the manufacturer’s protocol. For Western blotting and mass spectrometry, beads were resuspended in the SDS sample buffer, boiled for 5 min, and loaded onto an 8% acrylamide gel.

### Mass spectrometry

The post-IP samples were subjected to SDS-PAGE and the subsequent in-gel trypsin digestion as described (Lin et al., 2003). The peptides were dried using the CentriVap Benchtop Vacuum Concentrator (Labconco Corp.), dissolved in 0.1% trifluoroacetic acid (TFA), and desalted with C18 Ziptips (Millipore) according to the manufacturer’s protocol. NanoLC−nanoESI-MS/MS identification was performed on an Ultimate system 3000 nanoLC system (ThermoFisher Scientific) connected to the Orbitrap Fusion Lumos mass spectrometer equipped with NanoSpray Flex ion source (ThermoFisher Scientific). Briefly, peptide mixtures conditioned in 0.1% formic acid were injected into the HPLC instrument and enriched on a C18 Acclaim PepMap NanoLC reverse phase column of 25 cm length, 75 μm internal diameter and 2 μm particle size with a pore of 100 Å (ThermoScientific Scientific). The mobile phase consisted of aqueous solvent (A) 0.1% formic acid, and organic solvent (B) 0.1% formic acid in acetonitrile. Peptides were separated by 90 min of the segmented gradient from 2% to 40% solvent B at a flow rate of 300 nL/min. The mass spectra were acquired using one full MS scan followed by data-dependent MS/MS of the most intense ions in 3 sec. Full MS scans were recorded with an automatic gain control (AGC) target set at 500,000 ions and a resolution of 120,000 at m/z=200. For MS/MS spectra, selected parent ions within a 1.4 Da isolation window were fragmented by high-energy collision activated dissociation (HCD) with normalized collision energies of 32 eV. HCD-MS/MS with a resolution of 15,000, AGC at 50,000, and potential charge status of 2+ to 7+ were performed. A dynamic exclusion duration for the selection of parent ions was set at 180 sec with a repeat count. Spectra were collected by Xcalibur tool version 4.1 (ThermoScientific Scientific). The resulting RAW files were directly processed and analyzed by using MaxQuant software version 1.6.0.16. MS/MS peaks were searched in the Andromeda peptide search engine against the forward and reverse (as a decoy database) of reviewed UniProtKB/Swiss-Prot human proteome database. The common contaminants list attached in MaxQuant was also provided during the search (Tyanova et al., 2016). Peptides with minimum of 6 amino acids and maximum trypsin missed cleavages of 3 were considered. Variable modifications including methionine oxidation, N-terminal acetylation, Serine/Threonine/Tyrosine phosphorylation, and Asparagine/Glutamine deamidation, and the fixed modification of carbamidomethyl cysteine were applied. Initial parent peptide and fragment mass tolerance were set to 4.5 and 20 ppm, respectively. False discovery rate (FDR) filtration of peptide-spectrum match and protein levels were applied at 0.05 and 0.01, respectively. Finally, proteins identified as reverse, common contaminations or matched with only one unique peptide were excluded.

### Western Blotting

Cells were lysed in cold RIPA buffer supplemented with protease inhibitors (Merck, 539137) and phosphatase inhibitors (GoalBio, HC100-008). Equal amounts of proteins were resolved by 8% and 10% SDS-PAGE, transferred onto PVDF membranes, and blocked in a blocking buffer (Visual protein, BP01-1L) for 1 hour at room temperature. Primary antibodies were diluted in the blocking buffer at 1:1000. Secondary antibodies—anti-mouse or anti-rabbit-HRP (Bio-Rad)—were diluted at 1:5,000. Antibodies used in this study were listed in Supplemental Table 2. Blots were developed by using LuminataTM Crescendo Western HRP Substrate (Millipore, WBLUR0500) and images were taken in ChemiDoc® Touch imaging system. Raw densitometry for each antibody was normalized to GAPDH using Image Studio Lite. All quantification was performed in at least triplicate on original unmodified blots. One-way ANOVA with a Dunnett multiple comparison test was performed to compare the normalized signal for each antibody using GraphPad Prism 7. Error bars represent mean normalized signals ± SEM.

### Statistics

Statistical analysis of expression data was performed by R for Mac (Version 3.5.0) and GraphPad Prism 7 for Mac software. Patient demographics and survival information were gathered from database registry, electronic medical record and Taiwan Cancer Registry (for NTUH cohort). Cox proportional hazard modeling was used to examine the effect of expression levels of various fucosyltransferases on overall survival (time until death). The correlation between all members in the FUTs family were analyzed using Pearson correlation coefficients. Statistical analysis of the experimental data, including migration/invasion assays, drug treatments, etc, were performed using Mann-Whitney test or one-way ANOVA with Dunnett’s test. Data are presented as mean ± standard errors. *p* value < 0.05 is considered statistically significant.

### Study approval

The study on primary tumor tissues from lung cancer patients was approved by the Institutional Review Board (IRB) of National Taiwan University Hospital (Approval No.: 201701010RINB). Written informed consent was obtained from all participants prior to inclusion in the study. Every participant was assigned a number and the data were analyzed at the group level for patient de-identification. All mice experiments were performed in compliance with an experimental protocol (20160126) approved by the NTU College of Medicine Institutional Animal Care and Use Committee (IACUC).

### Data availability

All RNA-seq data are available at the National Center for Biotechnology Information Gene Expression Omnibus (https://www.ncbi.nlm.nih.gov/geo) under the accession number GSE120622. All proteomics data are available at the MassIVE repository (https://massive.ucsd.edu/ProteoSAFe/static/massive.jsp) under the accession number: MSV000083028

## Author contributions

Conceptualization, H.-C.T. and C.-J.Y.; Methodology; Y.-H.J. and H.-H.L.; Software and Formal Analysis, Y.-W.C., S.-Y.L., G.A.O, and R.R.W.; Resources, H.-C.T., J.-Y.S. and C.-J.Y.; Investigation, H.-H.L., Y.-H.J., H.-H.H., Z.-C.H., and Y.-J.H.; Writing – Original Draft, H.-H.L. and H.-C.T.; Writing – Review & Editing, H.-C.T. and C.-J.Y.; Funding Acquisition, H.-C.T. and C.-J.Y.; Supervision, H.-C.T. and C.-J.Y.

## Acknowledgments

We acknowledge the service provided by the Flow Cytometric Analyzing and Sorting Core of the First Core Laboratory and Laboratory Animal Center of National Taiwan University College of Medicine, Instrumentation Center, National Taiwan University, as well as Confocal Microscopy Core Facility, Institute of Biomedical Sciences, Academia Sinica. We also thank Ms. Min-Chuan Hsu and Ms. Pei-Chun Tsai for their administrative assistance. This study was supported by Ministry of Science and Technology (MOST 107-2314-B-002 −241, MOST 105-2628-B-002 −040), and National Health Research Institutes (NHRI-EX106-10610BC).

